# Improving early epidemiological assessment of emerging *Aedes*-transmitted epidemics using historical data

**DOI:** 10.1101/300954

**Authors:** Julien Riou, Chiara Poletto, Pierre-Yves Boëlle

## Abstract

Model-based epidemiological assessment is useful to support decision-making at the beginning of an emerging *Aedes*-transmitted outbreak. However, early forecasts are generally unreliable as little information is available in the first few incidence data points. Here, we show how past *Aedes*-transmitted epidemics help improve these predictions. The approach was applied to the 2015-2017 Zika virus epidemics in three islands of the French West Indies, with historical data including other *Aedes*-transmitted diseases (Chikungunya and Zika) in the same and other locations. Hierarchical models were used to build informative *a priori* distributions on the reproduction ratio and the reporting rates. The accuracy and sharpness of forecasts improved substantially when these *a priori* distributions were used in models for prediction. For example, early forecasts of final epidemic size obtained without historical information were 3.3 times too high on average (range: 0.2 to 5.8) with respect to the eventual size, but were far closer (1.1 times the real value on average, range: 0.4 to 1.5) using information on past CHIKV epidemics in the same places. Likewise, the 97.5% upper bound for maximal incidence was 15.3 times (range: 2.0 to 63.1) the actual peak incidence, and became much sharper at 2.4 times (range: 1.3 to 3.9) the actual peak incidence with informative *a priori* distributions. Improvements were more limited for the date of peak incidence and the total duration of the epidemic. The framework can adapt to all forecasting models at the early stages of emerging *Aedes*-transmitted outbreaks.

## Introduction

Model-based assessments must be done in real time for emerging outbreaks: this was the case in recent years for MERS-CoV in the Middle East [1, 2, 3], Ebola virus in West Africa[4, 5, 6, 7, 8, 9, 10], chikungunya virus (CHIKV) [11] and Zika virus (ZIKV) [12, 13, 14] in the Americas. These analyses often focused on transmissibility and reproduction numbers rather than on forecasting the future impact of the epidemic. Indeed, forecasting is difficult before the epidemic reaches its peak, all the more when information on natural history, transmissibility and under-reporting is limited[15]. Yet, it is precisely at the beginning of an outbreak that forecasts would help public health authorities decide on the best strategies for control or mitigation.

Several methods have been used to make epidemic predictions, including exponential growth models [5, 16], sigmoid-based extrapolations [17], SIR-type models [18] and more realistic model accounting for spatial and population structure [19]. But in addition to specifying a model, selecting good parameter values is also essential to obtain good predictions. In models for directly transmitted diseases, this can come from realistic demographic and behavioral characteristics, for example the contact frequency between individuals [20], mobility patterns [21, 22, 23, 24], and from clinical and epidemiological characteristics like the duration of the serial interval [25]. Such information is less easily available and more limited for mosquito-transmitted diseases [26]. However, outbreaks of the same disease or diseases with similar routes of transmission may have occurred in the same or similar locations, so that relevant information may be recovered from the analysis of past outbreaks.

Here, we show that past outbreaks of *Aedes*-transmitted diseases can substantially improve the epidemiological assessment of diseases transmitted by the same vector from early surveillance data. We introduce a hierarchical statistical model to analyze and extract information from historical data and obtain *a priori* distributions for required epidemiological parameters [27]. The method is illustrated with ZIKV outbreaks in the French West Indies between December, 2015 and February, 2017, using historical data regarding CHIKV and ZIKV epidemics in French Polynesia and the French West Indies between 2013 and early 2015. We assess the improvement in predictability of several operational indicators using different choices of *a priori* distribution and according to epidemic progress.

## Methods

### Data

Surveillance data on the 2015-2017 ZIKV epidemics in Guadeloupe, Martinique and Saint-Martin was collected by local sentinel networks of general practitioners and reported weekly by the local health authorities (Fig. 1) [28]. Cases of ZIKV infection were defined as “a rash with or without fever and at least two signs among conjunctivitis, arthralgia or edema”. We obtained numbers of suspected cases by week for each island, extrapolated from the number of active sentinel sites (dataset **𝓓**1). In the West Indies, local health authorities described the situation as “epidemic” when incidence was larger than 1 per 2,000 population per week (i.e. 200 cases in Guadeloupe and Martinique [29], and 20 cases in Saint-Martin). Following this description, we defined the “*S*”(-tart) of the epidemic as the first week above this threshold, the “*P*”(-eak) date when incidence was the highest, and the “*E*”(-nd) of the epidemic as the third consecutive week below the threshold (to ascertain the downwards trend). The time interval from “*S*” to “*E*” corresponds to the period of “high epidemic activity”.

**Fig 1.**
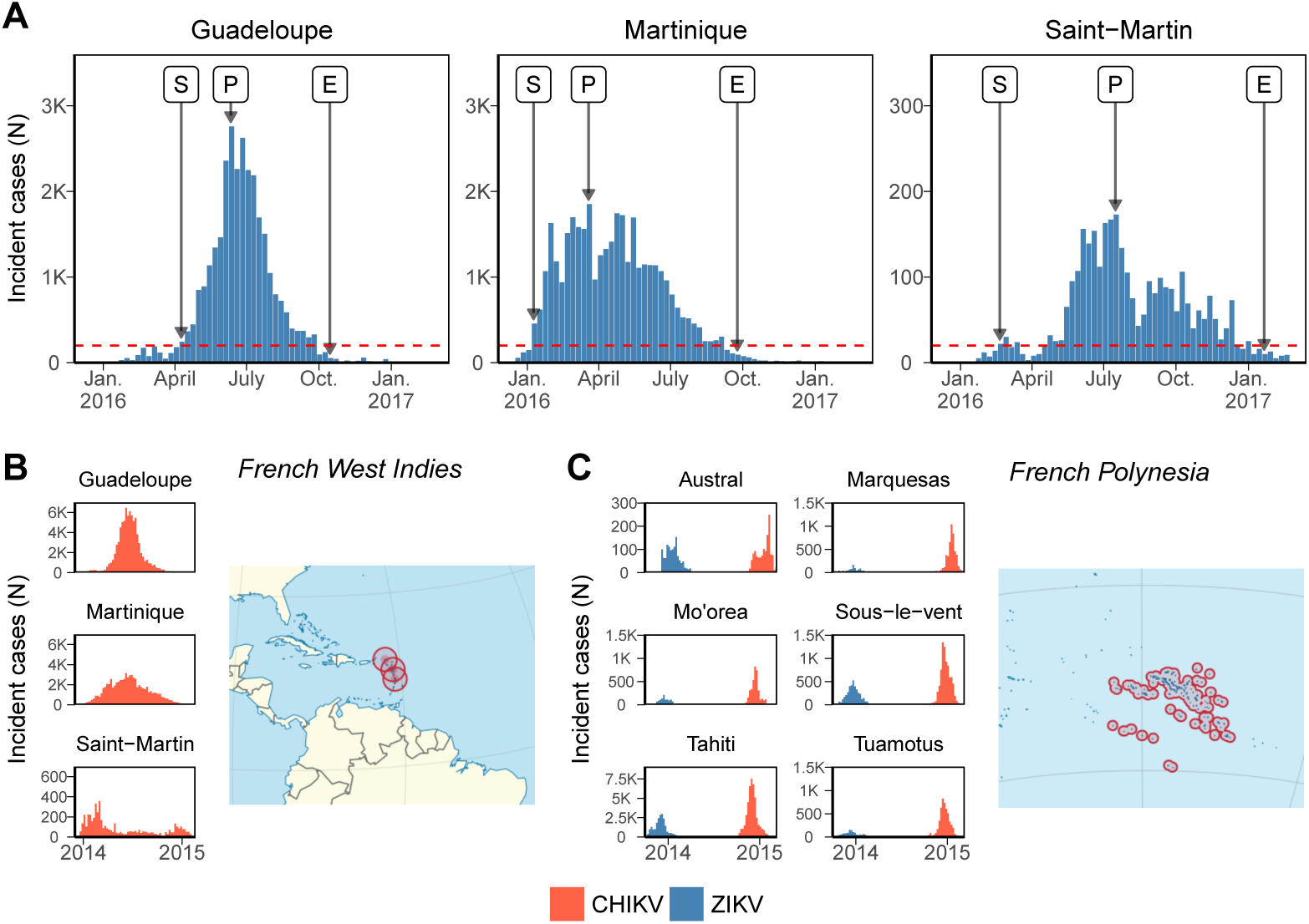
(A) Weekly number of Zika virus (ZIKV) cases reported by the surveillance systems in the French West Indies during 2016-2017 (dataset **𝓓**1). The dotted line shows the threshold defining high epidemic activity, “*S*” Mid “*E*” mark the start and the end of the period of high epidemic activity and “*P*” marks the date of peak incidence. (B-C) Weekly incidence (per 1,000 population) during the epidemics of chikungunya virus (CHIKV) in the French West Indies in 2013-2015 (dataset **𝓓**2) and of ZIKV then CHIKV in French Polynesia in 2013-2015 (dataset **𝓓**3).

We then analyzed historical data on the spread of emerging *Aedes*-transmitted diseases in similar locations. CHIKV epidemics occured in the same three islands during 2013-2015. Both diseases were transmitted by the same vector (*Aedes aegypti*), circulated in the same immunologically naive populations within a period of two years, had the same kind of clinical signs (i.e., fever, rash and arthralgia) and were reported by the same surveillance system. Surveillance data on CHIKV epidemics in the French West Indies was available from local health authorities (dataset **𝓓**2) [30]. Finally, we also selected the ZIKV and CHIKV epidemics that occured in six islands or archipelagoes of French Polynesia between 2013 and 2017, as they provided information on the differences between the two diseases. Surveillance data regarding the outbreaks in French Polynesia (dataset **𝓓**3) was collected following similar methods as in the French West Indies [31, 32].

### Epidemic model

The ZIKV outbreaks in Guadeloupe, Martinique and Saint-Martin were modelled separately using a dynamic discrete-time SIR model within a Bayesian framework. Briefly, the two main components of the model were: (i) a mechanistic reconstruction of the distribution of the serial interval of the disease (the time interval between disease onset in a primary case and a secondary case) that allows bypassing vector compartments; (ii) a transmission model for the generation of observed secondary cases in the human host. The generation time distribution was reconstructed by estimating the durations of each part of the infection cycle using disease- and mosquito-specific data from the literature, and assuming a fixed local temperature of 28°*C*, as described in more detail in the supplementary appendix. This led to gamma distributions with mean 2.5 weeks (standard deviation: 0.7) for ZIKV and with mean 1.6 weeks (sd: 0.6) for CHIKV.

Then, we linked weekly observed incidence *O_t_*,*_X_* to past incidence with:

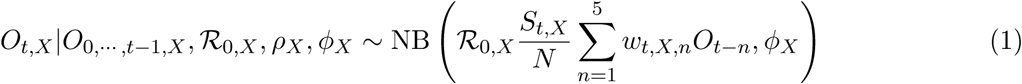

where subscript *X* refers to disease (*X* = *C* for CHIKV or *X* = *Z* for ZIKV), 𝓡_0,*X*_ is the basic reproduction number, *N* the population size, and 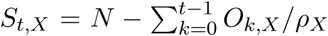 the number of individuals susceptible to infection at time *t* where *ρ_X_* is the reporting rate. The term 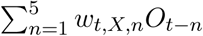 accounts for exposure to infection at time *t*, where *w_k_* is the discretized serial interval distribution. The variance is computed as the mean divided by the overdispersion parameter *ϕ_X_*. The model was implemented in Stan version 2.15.1 – R version 3.4.0 [33, 34, 35]. More details regarding the epidemic model and the Stan code are available in the supplementary appendix.

### Informative and non-informative prior distributions

To analyze Zika epidemics, the reproduction ratio 𝓡_0,*Z*_, the reporting rate *ρ_Z_* and the overdispersion parameter *ϕ_Z_* must be estimated. For *ϕ_Z_*, we used a non-informative prior in all cases [36]. For 𝓡_0,*Z*_ and *ρ_Z_*, we designed three different prior distributions, labelled as “non-informative” (NI), “regional” (R), or “local” (L) and described below.

The NI prior distributions expressed vague characteristics of the parameters: 𝓡_0,*Z*_ will be positive and likely not greater than 20; and *ρ_Z_* will range between 0 and 1.

The R and L priors were derived from the analysis of datasets **𝓓**2 and **𝓓**3 in three steps. We first analysed jointly the three CHIKV epidemics in dataset **𝓓**2 using model 1, introducing a two-level hierarchical structure for the island-specific parameters: an island-specific reproduction number 𝓡_0,*C*,*i*_ sampled from a top-level regional distribution 𝓝(*μ*_𝓡0,*C*_,*σ*_𝓡0,*C*_), and similarly logit(*ρC*,*i*) ∼ 𝓝(*μ_ρC_*,*σ_ρC_*) for the logit of the reporting rate. We obtained the posterior distributions of these parameters for the CHIKV outbreaks in each island *π*(𝓡_0,*C*,*i*, *ρC*,*i*_|**𝓓**2), as well as that of the hyperparameters *π*(*μ*_𝓡0,*C*_,*σ*_𝓡0,*C*_,*μ*_*ρ*,*C*_,*σ*_*ρ*_,*C*|**𝓓**2).

We then analysed dataset **𝓓**3 using the same hierarchical structure as above and introducing relative transmissibility and reporting of ZIKV with respect to CHIKV as 𝓡_0,*Z*_ = *β*_𝓡0_𝓡_0,*C*_ and *ρZ* = *β*_*ρ*_*ρC.* We thus used French Polynesia data to estimate the relative transmissibility *π*(*β_ρ_*|**𝓓**3) and the relative reporting rate *π*(*β_ρ_*|**𝓓**3) of ZIKV with respect to CHIKV in the French West Indies. In a third step, these posterior distributions were combined to obtain the R and L prior probability distributions as described in Table 1.

**Table 1.**
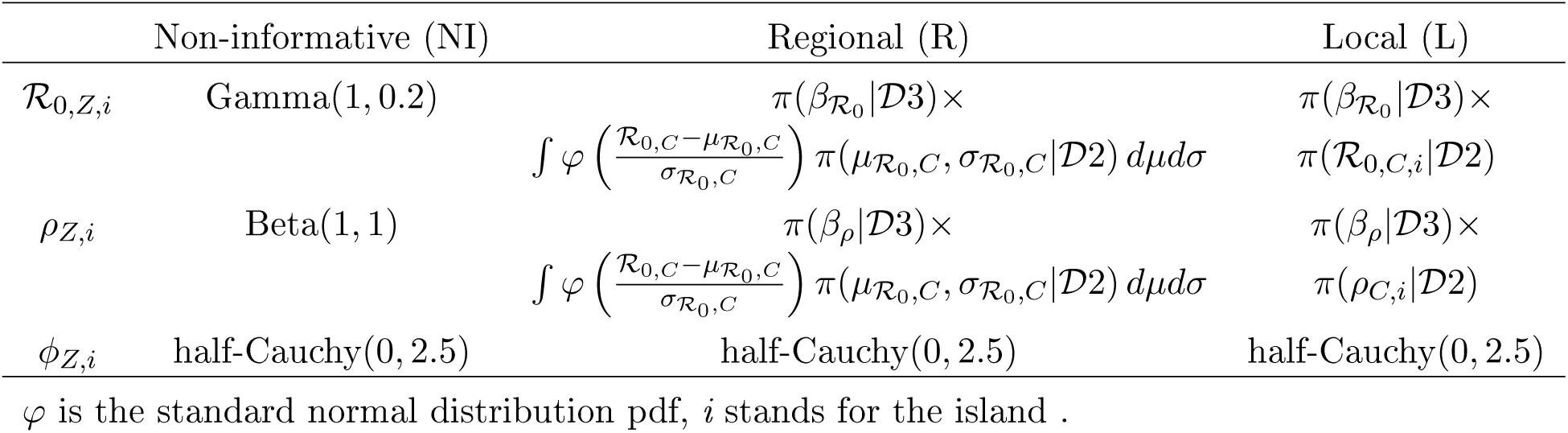
Prior distributions choice and design for modelling a Zika virus outbreak in the French West Indies.

The R and L priors differed in how much island-specific information they contained, the R priors being altogether less informative than the L priors. More precisely, the L priors used the bottom level in the hierarchical description, actually combining island-specific distributions for transmission
and reporting of CHIKV with relative ratios *β*_𝓡_0__ and *β_ρ_* of ZIKV to CHIKV. The R priors, on the contrary, were based on the top-level distributions in the hierarchical description, and can be interpreted as providing information for a “typical” island of the French West Indies rather than for a specific island.

Alternative prior distributions were considered in the sensitivity analysis and results are reported in the supplementary appendix.

### Fitting and predicting ZIKV epidemics

We fitted model 1 to ZIKV data separately in Martinique, Guadeloupe and Saint-Martin using the *K* first weeks of data (varying *K* from 5 to the number of weeks in the epidemic) and each *a priori* distribution (NI, R or L), to obtain posterior distributions for parameters 𝓡_0,*Z*_ *ρZ* and *ϕ_Z_* for every combination of island, *K* and choice of prior.

Then, using each set of posterior distributions, the epidemics were simulated forward from week *K* + 1 for two years using a stochastic version of the model described in equation 1. We used 16,000 replicates to compute the predictive distribution of the weekly number of future incident cases and a trajectory-wise 95% prediction band [37]. Using these simulated trajectories, we also computed the predictive distributions of four indicators of operational interest, for direct comparison with observed values:

- the final epidemic size, defined as the average total incidence across all simulated trajectories;
- the peak incidence, defined as the maximal value of the upper bound of the trajectory-wise 95% prediction band, as it corresponds to the capacity needed to ensure continuity of care [38];
- the date of peak incidence, defined as the average date of peak incidence across all trajectories;
- the epidemic duration, defined as the average duration between dates “*S*” and “*E*” across all trajectories.

The predictive distributions were compared using two measures of forecasting quality: (i) accuracy, i.e. the root-mean-square difference between predicted and observed values, and (ii) sharpness, i.e. the mean width of the 95% prediction band [39]. In order to improve clarity, these values were multiplied by –1 so that a higher value means a better accuracy or sharpness.

## Results

### ZIKV epidemics in the French West Indies

The timecourse of the ZIKV epidemics in Guadeloupe, Martinique and Saint-Martin between December, 2015 and February, 2017 differed markedly: the initial growth was early and sudden in Martinique, while it was delayed in Guadeloupe and Saint-Martin, starting only after four months of low-level transmission (Fig. 1). The epidemic showed a sharp peak in Guadeloupe, reaching a maximal weekly incidence of 6.9 cases per 1,000 inhabitants 9 weeks after the start of the period of high epidemic activity. In Martinique and Saint-Martin, weekly incidence reached a maximum of 4.8 cases per 1,000 inhabitants after a period of 10 and 21 weeks, respectively. Conversely, the period of high epidemic activity was longer in Martinique and Saint-Martin (37 and 48 weeks, respectively), than in Guadeloupe (27 weeks). In the end, a total of about 37,000 cases were observed in Martinique (97 cases per 1,000 inhabitants), more than in Saint-Martin (90 cases per 1,000 inhabitants) and Guadeloupe (77 cases per 1,000 inhabitants).

### Prior information from past epidemics

The CHIKV epidemics observed in the same three islands of the French West Indies during 2013-2015 are shown in Fig. 1B, and the CHIKV and ZIKV epidemics observed in French Polynesia during 2013-2015 in Fig. 1C. The *a priori* distributions on the reproduction ratio and reporting rates for the ZIKV epidemics in the French West Indies defining the NI, R or L approaches are shown in Fig. 2. The R priors were wide, with 95% credible intervals between 0.5 and 2.5 for 𝓡_0,*Z*_ and between 0 and 0.30 for *ρZ.* On the contrary, the more specific L priors on 𝓡_0,*Z*_ were highly concentrated around 1.5 in Guadeloupe and 1.3 in Martinique, and ranged between 1.0 and 1.8 in Saint-Martin. Likewise, the island-specific priors on *ρZ* carried more information that their regional counterpart, peaking around 0.19 in Guadeloupe, and covering wider intervals in Martinique (0.20-0.40) and Saint-Martin (0.03-0.39).

**Fig 2.**
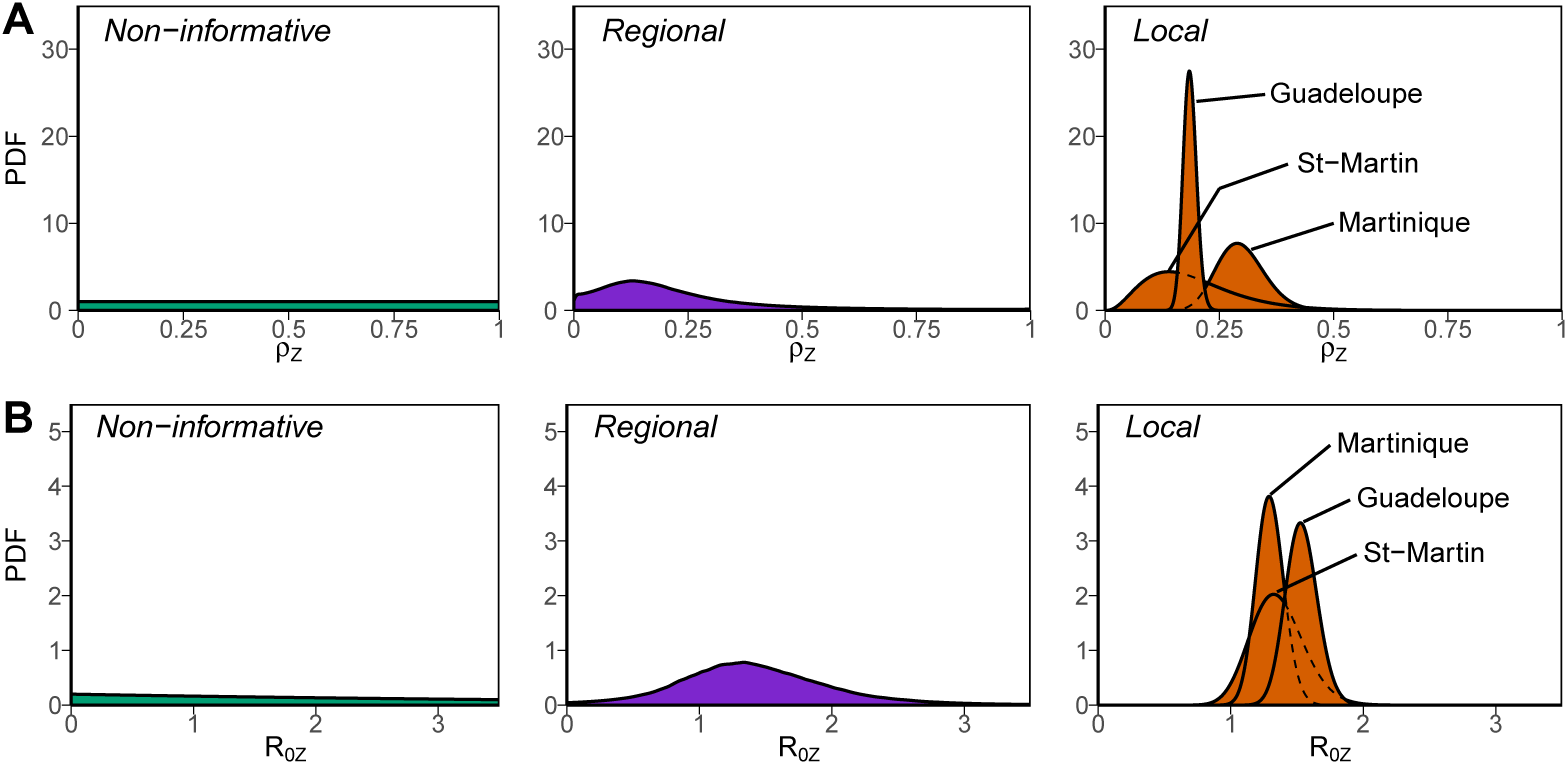
*A priori* distributions considered for the reporting rate *ρZ* (panel A) and the basic reproduction number 𝓡_0,*Z*_ (panel B) during the Zika virus epidemics in the French West Indies: non-informative, regional and island-specific.

### Epidemiological parameters and predictive distribution of future incidence

Fig. 3 shows the future course of the ZIKV epidemics predicted using data available two weeks after date “*S*” in each island, for the three choices of prior distributions. At this week, predictions with the NI priors were largely off-target, overestimating the future magnitude of the epidemic in all three islands. Using the R priors reduced the gap between forecasts and future observation. Major improvements in both *accuracy* and *sharpness* were obtained only with L priors. These results were typical of the initial phase of the epidemics, as shown in Fig. 4. Overall, the quality of the forecasts improved as *K* increases in all three islands and also as prior distributions brought more specific information. Results of the sensitivity analysis show that L priors defined here outperformed or performed at least equivalently to the alternative priors definition tested (supplementary appendix).

**Fig 3.**
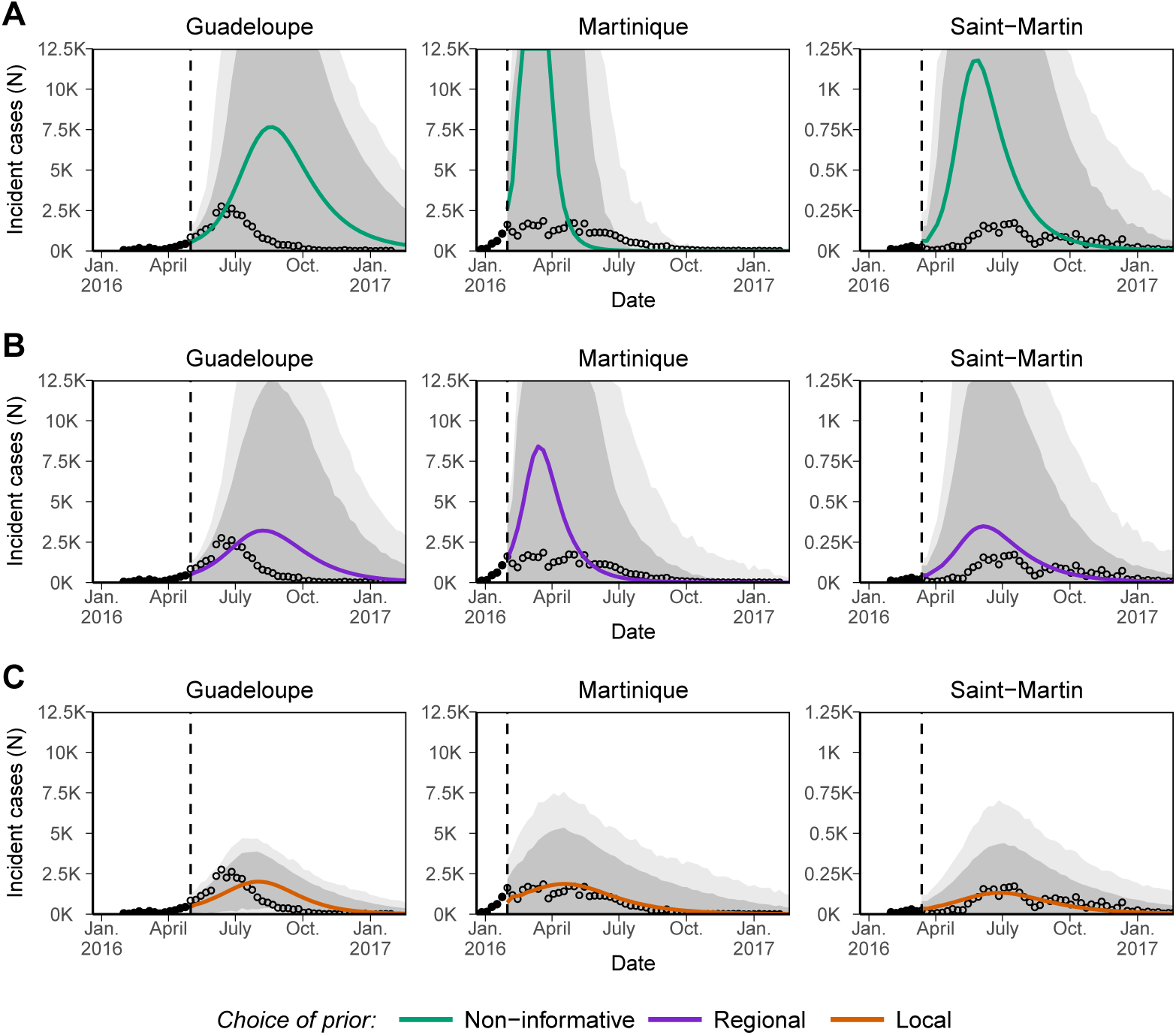
Predictive distribution of weekly incidence of Zika virus infections in Guadeloupe, Martinique and Saint-Martin using on either non-informative (NI, panel A), informative regional (R, panel B) or informative local (L, panel C) priors, and calibrated using data available up to the vertical dashed line (here chosen two weeks after date “*S*”). Continuous lines correspond to mean prediction of future incidence, dark and light grey areas to 50% and 95% prediction intervals, respectively, and circles to observed incidence.

**Fig 4.**
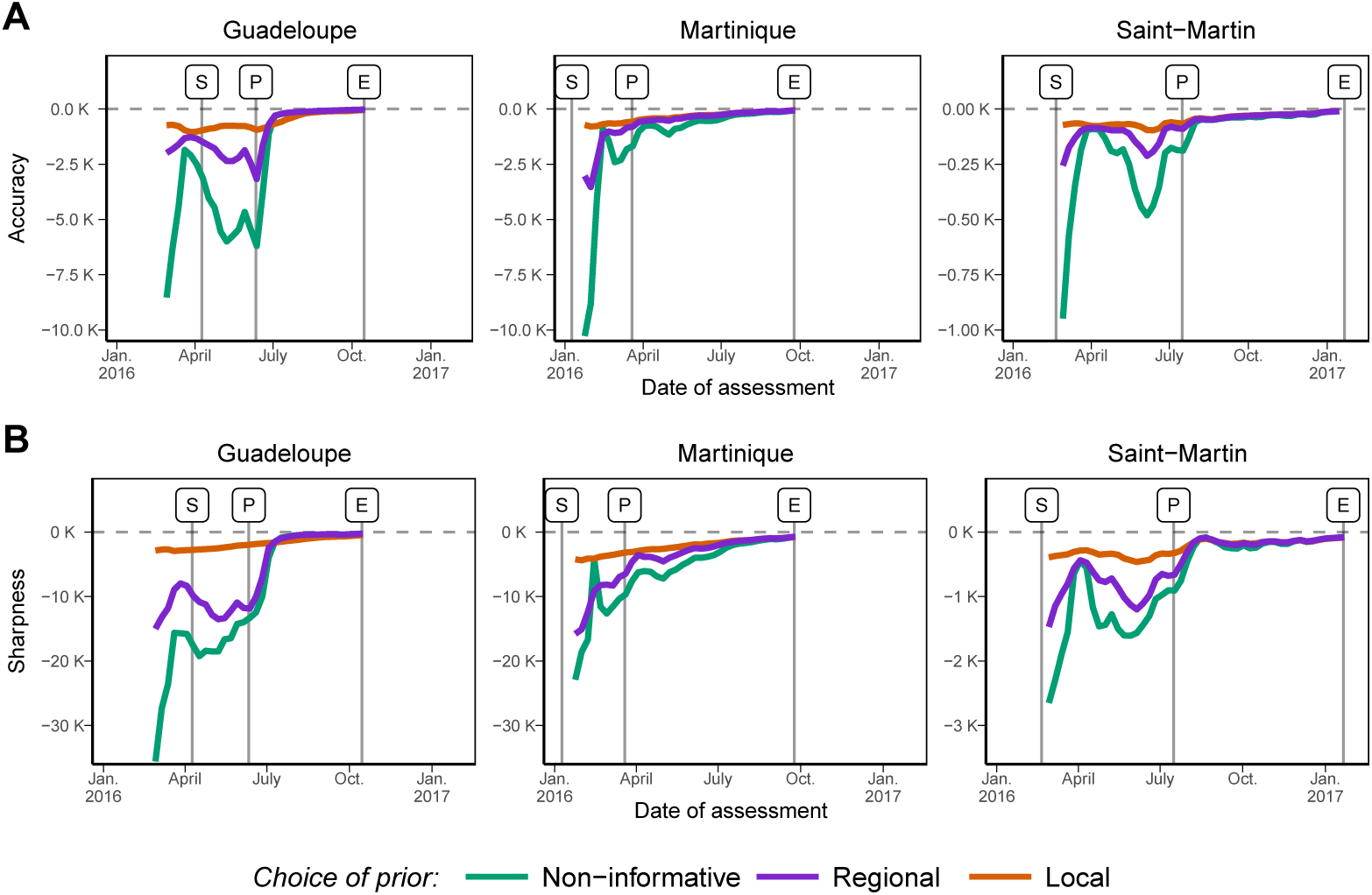
Accuracy (panel A, values closer to zero indicate better accuracy) and sharpness (panel B, values closer to zero indicate better sharpness) of the predictive distribution of future incidence based on epidemiological assessments conducted each week. Colours correspond to different *a priori* distributions on the parameters: non-informative priors or informative priors based on historical data, either considered at the regional or the local level.

The posterior distributions of the parameters built up differently as data accrued for 𝓡_0,*Z*_ and *ρZ*. For all three choices of prior distributions, the posterior distributions of 𝓡_0,*Z*_ quickly overlaid after a few points of incidence data were observed (Fig. 5A). In sharp contrast, the posterior distributions of *ρZ* could remain affected by the choice of prior distributions (Fig. 5B). In Martinique and Saint-Martin, *ρZ* remained essentially unidentified with the NI priors for the entire duration of the epidemic, with 95% credible intervals ranging approximately from 20 to 80%, even though the posterior mean was close to the estimates obtained with the more informative priors (around 25% at the end). Informative priors allowed for a more precise estimation of *ρZ*, and this remained the case over the whole course of the epidemics. In Guadeloupe, all posterior distributions for *ρZ* were similar after the peak, irrespective of the choice of priors.

**Fig 5.**
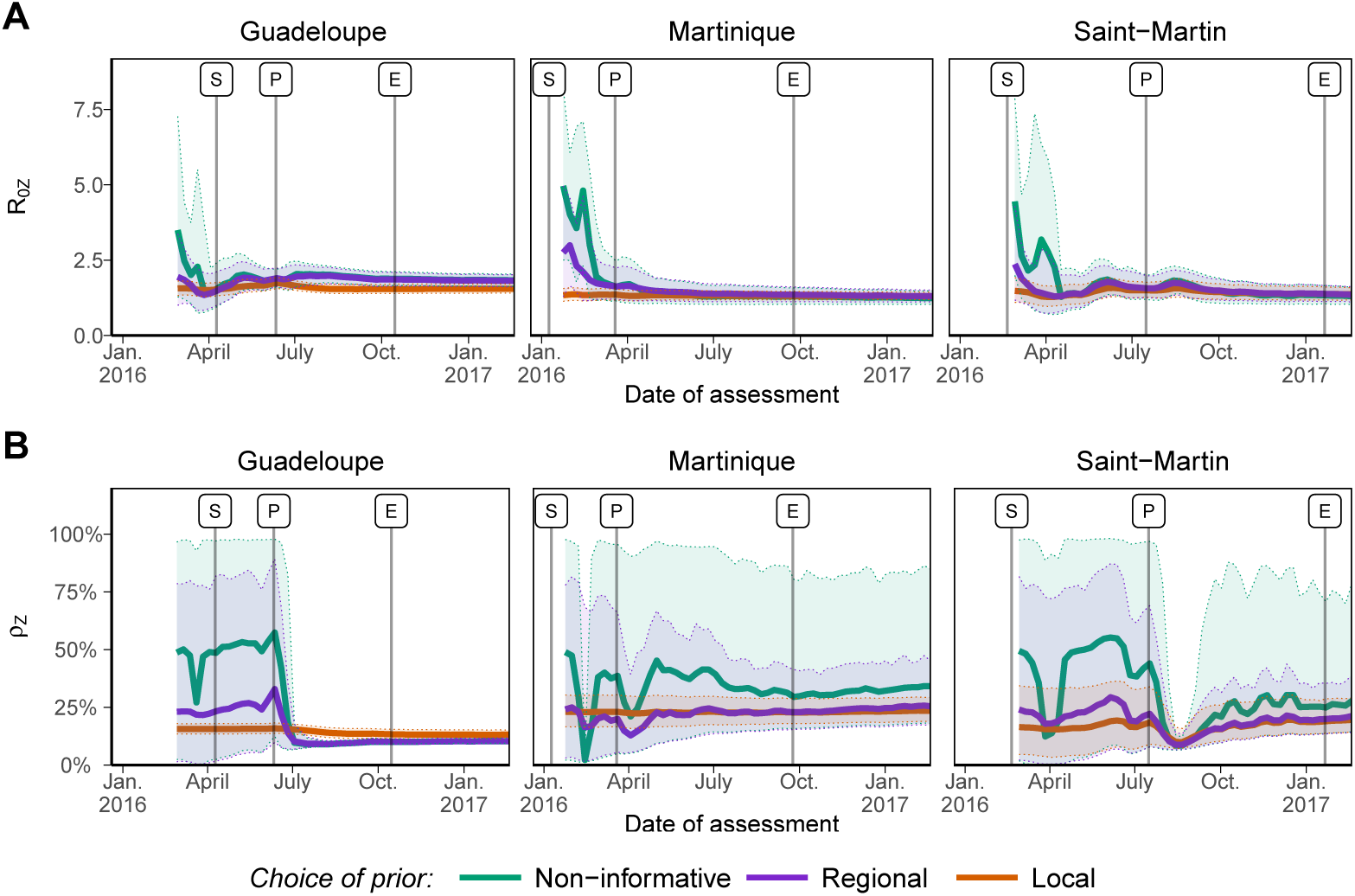
Posterior distributions (mean and 95% credible intervals) of the basic reproduction number 𝓡_0,*Z*_ (panel A) and the reporting rate *ρZ* (panel B) throughout the ZIKV epidemics of the French West Indies. Colours correspond to different *a priori* distributions on the parameters: non-informative priors or informative priors based on historical data considered either at the regional or the local level.

### Operational indicators

The forecasts of the four indicators of operational interest produced before peak incidence were contrasted. With NI priors, forecasts of total epidemic size overestimated the final counts by on average 3.3 times (range: 0.2 to 5.8) (Fig. 6A), with substantial variations in the forecasts from one week to the next. For instance in Martinique, projections varied from 105,000 total observed cases (95% prediction interval [95%PI]: 5,300-340,000) on February 7th to 8,100 (95% PI: 5,700-13,700) on February 14th. For the same indicator, forecasts using the R priors were only 1.7 times too high (range: 0.4 to 3.3) and those produced using L priors were only 1.1 times too high (range: 0.4 to 1.5). As a comparison, with the L priors on February 7th, the forecast of epidemic size was 47,200 (95%PI: 20,900-71,100) in Martinique, much closer to the final count of 37,400 observed cases at the end of the epidemic in this island.

**Fig 6.**
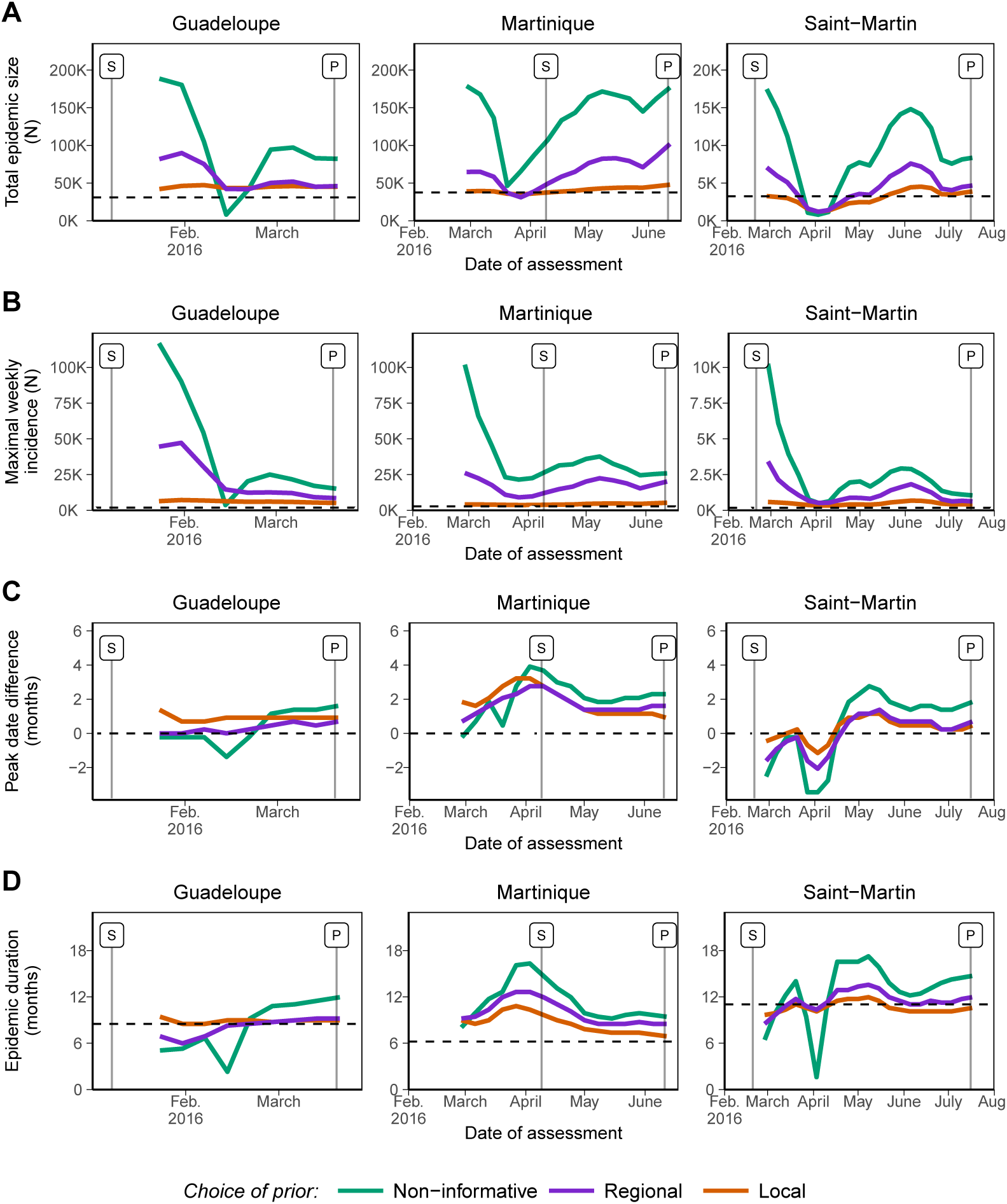
Early forecasts regarding four indicators of operational interest: (A) final epidemic size (total observed cases); (B) maximal weekly observed incidence; (C) date of peak incidence (difference with the date observed thereafter, in months); and (D) duration of the period of high epidemic activity (from date “*S* to date “*E*”, in months). The dashed lines represent the values observed after the end of the epidemic. Colours correspond to different *a priori* distributions on the parameters: non-informative priors or informative priors based on historical data, either considered at the regional or the local level.

Similarly, forecasts of maximal weekly (observed) incidence produced before date “*P*” were generally too large when using NI priors, on average 15.3 times higher than the actual maximal weekly incidence observed thereafter (range: 2.0 to 63.1) and with large fluctuations (Fig. 6B). Using informative prior distributions improved the forecasts, reducing the maximum predicted incidence to 7.5 times higher (range: 3.0 to 25.5) with the R priors and 2.4 times higher (range: 1.3 to 3.9) with the L priors. In all cases, forecasts of maximal incidence were never smaller than actual incidence.

Forecasting the dates of interest in the epidemics showed mixed results, with less differences depending on the choice of priors. The forecasts of the date of peak incidence were too late by on average 1.0 month (range: −3.4 to +3.9) using the NI priors, with large variations (Fig. 6C). Forecasts were only slightly better with the R priors (+0.7 months, range: −2.1 to +2.8) and the L priors (+0.9 months, range: −1.1 to +3.2), but sharper. Better forecasts were obtained for Martinique than for the other islands.

The forecasts of the total duration of the period of high epidemic activity were overestimated by a factor 1.3 (range: 0.1 to 2.6) with NI priors, again with high variability from week to week (Fig. 6D). Informative priors brought a small improvement, in particular regarding the stability of the forecasts over time, overestimating the actual duration by a factor 1.2 (range: 0.7 to 2.0) with R priors and by a factor 1.1 (range: 0.9 to 1.7) with L priors.

## Discussion

Obtaining reliable model-based forecasts in real time at the beginning of an epidemic is a difficult endeavour. It is however precisely during these periods that forecasts may have the most impact to guide interventions. Here, we compared several approaches to provide forecasts for ZIKV epidemics from an early point in a retrospective analysis of the outbreaks that occurred in the French West Indies in 2015-2017. We found that the accuracy and sharpness of the forecasts before peak incidence were substantially improved when *a priori* information based on historical data on past epidemics was used.

The three ZIKV outbreaks in the French West Indies provided an ideal situation to look for ways to improve prediction of *Aedes*-transmitted diseases using historical data. Indeed, CHIKV outbreaks had been observed in the same locations about two years before ZIKV, CHIKV is also transmitted by *Aedes* mosquitoes, and both viruses spread in a region where the populations were immunologically naive at first. Furthermore, all epidemics were observed by the same routinely operating GP-based surveillance networks, and the three locations benefit from a mature public health system, with easy access to medical consultation and individual means of protection. Last, pest control is done in routine, with additional intervention showing limited efficacy in this context [40, 41]. This motivated our decision to use constant parameters for transmsission and reporting over time in the modelling.

Having established the many similarities between ZIKV and CHIKV epidemics regarding transmission and reporting, we assumed that information could be transported (as defined in [42]) between diseases and between places. Bayesian approaches allow such transportation using informative *a priori* distributions on model parameters [27]. Historical data on CHIKV epidemics was therefore used to build informative *a priori* distributions on the two key parameters 𝓡_0,*Z*_ and *ρZ*. Hierarchical models are particularly adapted to this task, as they naturally pool information among several past epidemics and capture both within- and between-location variability [43]. We used separate hierarchical models to obtain information about (i) transmissibility and reporting during CHIKV outbreaks in the French West Indies and (ii) relative transmissibility and reporting between ZIKV and CHIKV in French Polynesia, rather than from a global joint model (as in [44]). This choice was made to show that prior information can be combined from separate sources in a modular way. We finally considered two versions of informative priors to capture different degrees of knowledge. The “(L)ocal” priors corresponded with estimates for the previous CHIKV epidemic in the same island, therefore including island-specific epidemic drivers, such as population structure and distribution, socio-economic circumstances and environmental conditions. On the other hand, the “(R)egional” priors encompassed the diversity of past observations within the region, making priors valid for a typical island of the West Indies, especially when no other epidemic has been observed previously.

Analyses conducted at the early stage of an epidemic using non-informative *a priori* distributions – as is often done – led to poor forecasts before the peak of incidence was reached, an observation already made in other studies [45]. Indeed, early forecasts of epidemic size were largely off-target and unstable, varying between 0.2 and 5.8 times the eventually observed total incidence. Worst case projections on maximal incidence were very imprecise, ranging between 2 and 63 times the eventually reached maximal weekly incidence. Using historical data led to a substantial increase of the quality of these forecasts from the very early stages of the epidemics. Using “local” priors, the ratio between forecasts and reality ranged between 0.4 and 1.5 for epidemic size and between 1.3 and 3.9 for maximal incidence. The less specific “regional” priors increased accuracy and sharpness as well, though to a lesser extent. However, the date of peak incidence and the date of the end of the period of high epidemic activity were only slightly improved by integrating historical data.

The posterior distributions of all forecasted quantities changed as more data was included (Fig. 4). The posterior estimates of 𝓡_0,*Z*_ were quickly similar and in good agreement with prior information. On the contrary, the reporting rate *ρZ* remained essentially unidentifiable until after the incidence peak in Guadeloupe, and to the end of the outbreak in Martinique and Saint-Martin. This suggests that prior information is essentially required for the reporting rate, a difficult to estimate quantity as already noted [15]. Sensitivity analysis support this result. Indeed, providing informative prior on p_z_ only leads to similar results as providing it for both *ρZ* and 𝓡_0,*Z*_ (supplementary appendix).

Predicting the future course of epidemics from an early point is increasingly seen as a problem of interest [46, 9, 8], and forecasting challenges have been set up for influenza [47], for Ebola [45] and for chikungunya [48]. Comparing and systematically evaluating models’ forecasting performances is still at the beginning. As of now, comparisons targeted the merits of different models including exponential growth models, sigmoid models, or mechanistic epidemic models [45]. Our work provides a complementary approach where information from past epidemics is combined using hierarchical models to inform on parameter ranges, thus increasing the reliability of early forecasts. It was applied here to a dynamic discrete-time SIR model that for its parsimony is well-adapted to real-time forecasting. Complex mechanistic models can provide a more realistic description of the epidemic, accounting, for instance, for heterogenous spatial distribution of individuals and mobility coupling – a relevant ingredient for describing epidemics in more extended spatial areas –, or vector population dynamics and its mixing with humans. Our framework could in principle be adapted to these more sophisticated models.

Only recently have hierarchical models been used for modelling multiple epidemics, for instance with the joint analysis of six smallpox epidemics [49], for transmissibility and duration of carriage in the analysis of multistrain pneumococcus carriage [50], or for forecasting seasonal influenza [51].

Other modelling papers specifically attempted to take advantage of the similarities between different *Aedes*-transmitted diseases, e.g., by estimating the risk of acquiring chikungunya from the prevalence of dengue [52] or by assesssing the spatio-temporal coherence of chikungunya, Zika and dengue [53]. Also, using informative priors to make up for the lack of information during the early stages of an epidemic has been done before. For instance, *a priori* information from the ZIKV epidemics in French Polynesia has been used to support the early forecasts of health-care requirements for the ZIKV epidemic in Martinique [38]. In this case, however, authors concluded that a prior built from an epidemic in a different location resulted in inaccurate predictions at the early stage. We found a similar result in a sensitivity analysis: the direct use of information from ZIKV in French Polynesia, or alternatively the direct use of information from CHIKV in the French West Indies, without adjusting for ZIKV, leads to poor overall forecasting quality compared to the L prior considered here (supplementary appendix).

This shows that the choice of appropriate historical data is the cornerstone of any such attempt. Yet little is known regarding the comparative epidemiology of diseases in the same or similar places and on the condition where transportability can be assumed. For influenza, it has been reported that the reproduction ratios in two successive flu pandemics (1889 and 1918) showed substantial correlation (*r* = 0.62) in the same US cities, even years apart [54]. For *Aedes*-transmitted diseases, comparisons of ZIKV and dengue virus outbreaks [55] and of ZIKV with CHIKV outbreaks [44] in the same locations have highlighted similarities in the epidemic dynamics. In any case, careful consideration of all the factors that may influence transmission and reporting is needed. For example, contrary to CHIKV, some cases of sexual transmission have been reported for ZIKV [56, 57]. Yet, no epidemics were seen in locations without enough *Aedes* mosquitoes, as for example in metropolitan France, despite the introductions of several hundreds ZIKV infected cases [58]. This justifies the use of CHIKV epidemic data to provide prior information on ZIKV in the epidemic context considered here. More generally, documenting, analyzing and comparing more systematically past epidemics [59] is necessary to provide the data required to derive prior information. In particular, informed with epidemiological records of the recent CHIKV and ZIKV epidemics, our approach could be applied to potential future emergences of other *Aedes*-transmitted diseases such as Mayaro virus [60], Ross River virus [61] or Usutu virus [62] once transportability is deemed plausible.

## Supporting information

**Supplementary appendix.** Detailed description of the models (including Stan code), additional results and sensitivity analyses.

**Animated fig. S1:** Forecasted mean weekly number of clinical cases from analyses conducted at every point of the epidemics (animated GIF file).

**Dataset 𝓓1**: Weekly number of reported cases of Zika virus infection in the French West Indies (2015-2017).

**Dataset 𝓓2**: Weekly number of reported cases of chikungunya virus infection in the French West Indies (2013-2015).

**Dataset 𝓓3**: Weekly number of reported cases of Zika virus and chikungunya virus infection in French Polynesia (2013-2015).

## Acknowledgments

We thank both the *CIRE Antilles*-*Guyane* and the *Centre d’Hygiène et de Salubrité Publique de Polynésie française* for collecting the data and making it publicly available.

